# Pan-cancer immunogenomic analyses reveal genotype-immunophenotype relationships and predictors of response to checkpoint blockade

**DOI:** 10.1101/056101

**Authors:** Pornpimol Charoentong, Francesca Finotello, Mihaela Angelova, Clemens Mayer, Mirjana Efremova, Dietmar Rieder, Hubert Hackl, Zlatko Trajanoski

## Abstract

The Cancer Genome Atlas revealed the genomic landscapes of common human cancers. In parallel, immunotherapy with checkpoint blockers is transforming the treatment of advanced cancers. As only a minority of the patients is responsive to checkpoint blockers, the identification of predictive markers and the mechanisms of resistance is a subject of intense research. To facilitate understanding of the tumor-immune cell interactions, we characterized the intratumoral immune landscapes and the cancer antigenomes from 20 solid cancers, and created The Cancer Immunome Atlas (http://tcia.at). Cellular characterization of the immune infiltrates revealed a role of cancer-germline antigens in spontaneous immunity and showed that tumor genotypes determine immunophenotypes and tumor escape mechanisms. Using machine learning we identified determinants of tumor immunogenicity and developed a scoring scheme for the quantification termed immunophenoscore. The immunophenoscore was superior predictor of response to anti-CTLA-4 and anti-PD-1 antibodies in two independent validation cohorts. Our findings and the developed resource may help informing cancer immunotherapy and facilitate the development of precision immune-oncology.

## INTRODUCTION

In the past decade, driven by technological advances and novel mechanistic insights we are witnessing two major advances in cancer research and treatment. First, the emergence of next-generation sequencing (NGS) technologies and comprehensive and coordinated efforts (e.g. The Cancer Genome Atlas (TCGA)) enabled the characterization of multi-dimensional maps of genomic changes in common cancers. And second, basic research in cancer immunology paved the way for the development and approval of checkpoint blockers. These drugs which augment T cell activity by blocking cytotoxic T lymphocyte antigen-4 (CTLA-4), programmed cell death protein 1 (PD-1), or PD-1 ligand (PD-L1), show remarkable clinical effects. Analysis of long-term data of patients who received anti-CTLA-4 antibodies in unresectable or metastatic melanoma shows a plateau in the survival curve after 3 years (Schadendorf et al., 2015), suggesting curative potential. Over and above, efficacy of anti-PD-1 antibodies has been shown not only in melanoma, but also in nine different tumor types (Wolchok, 2015). There is currently a rapid pace of development of checkpoint blockers evident from more than 150 clinical trials with monotherapies or combination therapies (Wolchok, 2015). However, there is disparity in response rates across and within tumor types, suggesting the existence of intrinsic immune resistance, as well as evidence for acquired immune resistance (Pitt et al., 2016).

With the development of the immunotherapies with checkpoint blockers as well as other immunotherapeutic strategies including therapeutic vaccines and engineered T cells (Schumacher and Schreiber, 2015), the tumor-immune cell interaction came into focus. The investigation of tumor-immune cell interaction poses considerable challenges, due to the evolving nature of this two ecosystems: the development of cancer, which can be seen as evolutionary process, and the immune system, with a number of innate and adaptive immune cell subpopulations, some of which show phenotypic plasticity and possess memory. Using genomic data and bioinformatics tools it is now possible to computationally dissect tumor-immune cell interactions (Hackl et al., 2016). These immunogenomic analyses can provide information on the two crucial characteristics of the tumor microenvironment: 1) composition and functional orientation of the infiltrated immune cells, and 2) expression of the cancer antigenome, i.e. the repertoire of tumor antigens including two classes of antigens: neoantigens which arise from somatic mutations, and cancer-germline antigens (CGA).

Previous studies have used genomic data from the TCGA to characterize neoantigens and their association with survival (Brown et al., 2014) or with cytolytic activity estimated using the expression of two genes (Rooney et al., 2015). However, these studies did not characterize the cellular composition of the intratumoral immune infiltrates. More recently, gene signature were used to analyze infiltration of B cells, T cells, and macrophages and the prognostic relevance of these subpopulations (Iglesia et al., 2016). Similarly, RNA expression data corrected for tumor purity was used to estimate infiltration of B cells, CD4^+^ T cells, CD8^+^ T cells, neutrophils, macrophages and dendritic cells. (Li et al., 2016). However, while such analyses of few major cell types are helpful for identifying clinical associations, higher resolution of the TIL landscape is required in order to dissect tumor-immune cell interactions, and identify prognostic and predictive markers. We have previously shown that the tumor infiltrates are composed of at least 28 different types (Bindea et al., 2013), some of which are generally associated with favorable prognosis, whereas others like regulatory T cells (Tregs) are immunosuppressive (deLeeuw et al., 2012). Over and above, we described the importance of memory and cytotoxic cells in controlling the growth and recurrence of tumors in colorectal cancer (CRC) (Galon et al., 2006) and showed that a prognostic marker termed immunoscore based on the quantification of these two subpopulations more accurately predicts survival than the standard TNM staging (Mlecnik et al., 2011). Thus, it is of utmost importance to provide a comprehensive view of the intratumoral immune landscape including memory cells, cytotoxic cells (CD8 ^+^ T cells, natural killer (NK) cells, and NK T (NKT) cells), as well as immunosuppressive cells (Tregs and myeloid-derived suppressor cells (MDSCs)).

We have recently developed an analytical strategy to characterize the cellular composition of the immune infiltrates and examined colorectal cancer (CRC) datasets from the TCGA (Angelova et al., 2015). Our approach is based on the use of metagenes, i.e. non-overlapping sets of genes that are representative for specific immune cell subpopulations and are neither expressed in CRC cell lines nor in normal tissue. The expression of these sets of metagenes is then used to analyze statistical enrichment using genes set enrichment analysis (GSEA). The advantage of the metagene approach is the robustness of the method due to two characteristics: 1) the use a set of genes instead of single genes that represent one immune subpopulation. The use of single genes as markers for immune subpopulations can be misleading since many genes are expressed in different cell types; and 2) the assessment of relative expression changes of a set of genes in relation to the expression of all other genes in a sample. Thus, the calculations are less sensitive to noise resulting from sample impurity or sample preparation compared to the deconvolution methods.

Here, we further developed our metagene approach by defining a set of pan-cancer metagenes for 28 immune cell subpopulations and scaled up the analyses to solid cancers. We carried out immunogenomic characterization of the TCGA data for 20 solid cancers with >8000 tumor samples, and provide for the first time comprehensive view of the cellular composition of the intratumoral immune infiltrates. Additionally, we derived also cancer antigens as well as genetic characteristics (tumor heterogeneity and clonality) for individual samples in order to enable integrative analyses of both, immune features as well as genetic features of the tumors. We then developed a web-accessible database TCIA (The Cancer Immunome Atlas) with the results of our analyses (<u>http://tcia.at</u>). To demonstrate the utility of the resource we carried out integrative analyses and revealed cellular profiles that were predictors of survival for distinct cancers and genotype-immunophenotype relationships. We further demonstrate the value of the TCIA by using machine learning approach to identify determinants of immunogenicity and propose a novel scoring scheme for solid cancers: the immunophenoscore. Validation with two cohorts treated with anti-CTLA-4 and anti-PD-1 antibodies showed that the immunophenoscore is a superior predictor of response to checkpoint blockers in patients with melanoma.

## RESULTS

### High-resolution genomic analyses of the tumor-immune interface

An overview of the strategy for the immunogenomic analyses and the used methods is shown in Figure 1a (for details see Computational Methods and Supplementary Table S1). We mined the TCGA data to extract the following information: 1) TILs, which were estimated from gene expression data using two approaches: GSEA and deconvolution, 2) cancer antigenomes comprising neoantigens and CGAs, 3) tumor heterogeneity, and 4) immunophenotypes, which were defined using TILs and predetermined sets of genes (MHC molecules, immunomostimulators, and immunoinhibitors).

**Figure 1.**
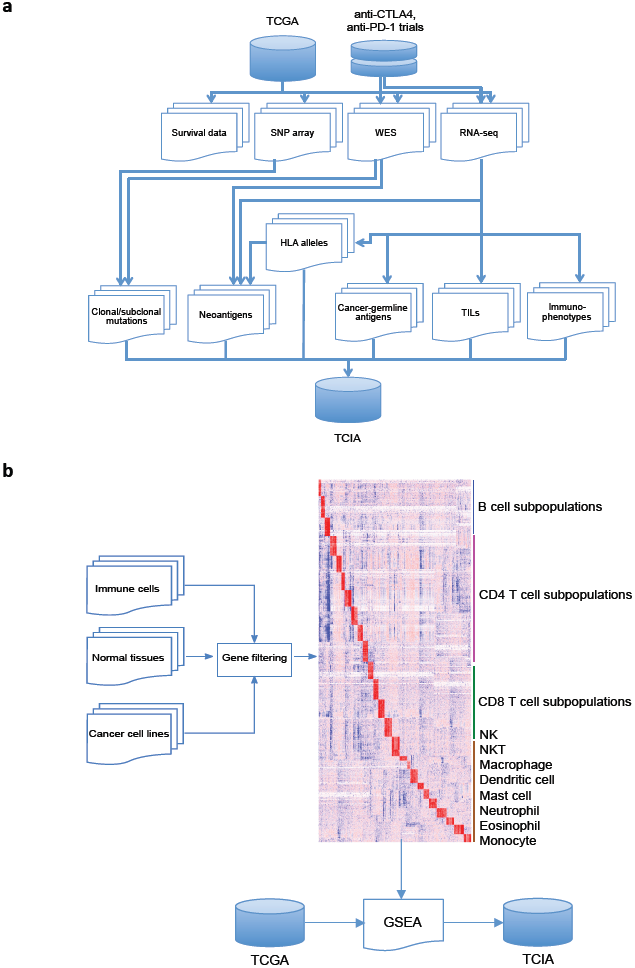
Strategy for pan-cancer immunogenomic analyses. (**a**) The scheme shows the immunogenomic analyses and the types of data used for the analyses. The results are deposited in a web-accessible database The Cancer Immunome Atlas (TCIA) (<u>http://tcia.at</u>). (**b**) Immune-related signatures are derived from expression profiles of purified immune cells, normal cells, and cancer cell lines, and used for the gene set enrichment analysis (GSEA) of the TCGA RNA-sequencing data.

To estimate TILs we first built a compendium of genes (1625) related to specific immune cells using gene expression profiles from sorted immune cell subpopulations from 37 studies comprising 366 microarrays (Figure 1b). A subset of these genes, which are representative for specific immune cell subpopulations and are neither expressed in cancer cell lines nor in normal tissue were then selected based on correlation analysis (782 in total). We then used gene set enrichment analysis (GSEA) (Subramanian et al., 2005) with this gene set to decompose cellular profiles from RNA sequencing data from bulk tissue for individual samples. Additionally, we used an independent method to identify fractions of immune subpopulations based on a deconvolution approach (CIBERSORT (Newman et al., 2015)). At the time of analysis, CIBERSORT was only optimized for microarray data, since the immune signature matrix was built from a microarray compendium. Therefore, we implemented a strategy to modify RNA-sequencing data from TCGA to be used as input for CIBERSORT and built a model to transform RNA-sequencing data to microarray data considering the TCGA samples for which microarray and RNA-sequencing data were available (n=550, see Computational methods). Despite the high correlation between microarray- and RNA-seq-based estimates for these samples (see Computational methods), a lower performance for samples where signature genes have a wider range of expression than the model cannot be excluded. Moreover, several important subpopulations were missing in the deconvolution method including activated and memory CD8^+^ T cell, and MDSCs. Therefore, we present here only the results from the GSEA method, which depicts a more comprehensive picture of the tumor-suppressive or tumor-promoting roles of TILs. However, we make both GSEA and deconvolution data available on the TCIA website because they are complementary (e.g. polarized macrophages are estimated only by the deconvolution method).

It has been previously shown that the tumor purity can be used in selecting genes informative for deconvolving TILs (Li et al., 2016). We therefore corrected the mutational load for tumor purity and analyzed associations of individual TIL subpopulations estimated using the GSEA and tumor purity (Supplementary Figure S1). As can be seen, there is a law to weak correlation suggesting that the GSEA method is robust to varying degrees of tumor purity.

To chart the antigenome for each sample we used RNA-sequencing data to derive expression levels of CGAs. For the identification of neoantigens we assembled an analytical pipeline as previously described (Angelova et al., 2015). Briefly, for each subject 4-digit HLA class I alleles were predicted from RNA-sequencing data. Peptides of 8-11 amino acids in length, covering the mutated region of the protein were then generated. The predicted HLA alleles and the mutated expressed peptides for each samples were used as input for the algorithm NetMHCpan (Nielsen et al., 2007) to estimate their binding affinities and predict neoantigens.

Tumor mutational heterogeneity adds another layer of complexity (Gerlinger et al., 2012): within a tumor, clones may be present which do not elicit T-cell responses against a given neoantigen and these cells may have selective advantage and outgrow other clones. We therefore estimated also tumor heterogeneity using exome sequencing data and SNP arrays data to calculate cancer cell fractions (CCF). Finally, driver genes were assigned from a recently published study (Rubio-Perez et al., 2015) whereas clonal and subclonal origins of the neoantigens were estimated as previously described (Landau et al., 2013).

Using this strategy and the analytical pipelines we developed, we analyzed TCGA data for 8243 samples and 20 solid tumors. Hereafter colon (COAD) and rectal (READ) cancers are considered as single entity (CRC). Additionally, available exome-seq and RNA-sequencing data from recent clinical studies using antibodies against CTLA-4 (Van Allen et al., 2015) and PD-1 (Hugo et al., 2016) were analyzed using the same procedures. In total, 1 PBytes of analyzed primary and processes data genomic data resulted in 50 Gbytes of structured immunogenomic data.

We then developed a web-accessible relational database TCIA (<u>http://tcia.at</u>) and deposited the results of the analyses including immune infiltrates calculated with both, GSEA and the deconvolution method, expression of predefined immune subsets, CGAs, HLA alleles, neoantigens, tumor heterogeneity, and clinical data. The database allows queries for the above features for user-defined cancers and subgroups of patients, as well as free download of the data for further analyses. Due to privacy concerns, access to HLA allele data is provided only to researchers who have access to the primary sequence data from the TCGA.

### Cellular characterization of immune infiltrates reveals prognostic cell types in distinct cancers

Using our GSEA strategy we estimated 28 subpopulations of TILs including major types related to adaptive immunity: activated T cells, central memory (Tcm), effector memory (Tem) CD4^+^ and CD8^+^ T cells, gamma delta T cells (Tγδ), T helper 1 (Th1), Th2, Th17, regulatory T cells (Treg), follicular helper T cells (Tfh), activated, immature, and memory B cells, as well as cell types related to innate immunity: macrophages, monocytes, mast cells, eosinophils, neutrophils, activated, plasmocytoid and immature dendritic cells (DCs), NK, natural killer T cells (NKT), and MDSCs.

The results of the cellular characterization of the immune infiltrates using GSEA showed heterogeneity across cancers and within individual cancer entities and associations of specific TIL subpopulations with survival (Figure 2a). The cellular profiles associated with survival differed between cancers. In general, the infiltration of many TIL subpopulations related to adaptive immunity was associated with good prognosis including activated CD8^+^ T cells, Tem and Tcm CD8^+^ cells, and Tem CD4^+^ cells, whereas MDSCs and Tregs were associated with bad prognosis. The most common cell type significantly associated with good prognosis was Tem CD8^+^ (15 out of 19 cancers) followed by activated CD8^+^ cells (10 out of 19 cancers) and Tem CD4^+^ (9 out of 19 cancers). These results were consistent with a number of previous studies we and others have published (Fridman et al., 2012). Cell types significantly associated with bad prognosis were MDSCs (13 out of 19 cancers) and Tregs (10 out of 19 cancers). The cellular profiles and the clinical information for specific cancers can be viewed at the web site and downloaded for further analyses.

## Mutational load and tissue context determine cellular composition of immune infiltrates

The vision of precision oncology implies patient specific genomic profiling and testing of targeted therapeutics on the subset of patients carrying the relevant genetic lesions, rather than on patients selected based on histologic tumor subtypes (Garraway and Lander, 2013). We asked what is consequence of this genome-centric view on the immune landscapes in solid cancers and analyzed the impact of the mutational load. The tumors were classified into three groups based on the mutational load: tumors with high mutational load (upper quartile), intermediate mutational load (two intermediate quartiles), and low mutational load (lower quartile) (Figure 2a). Tumors with high mutational load were enriched with activated T cells and Tem cells and were depleted with immunosuppressive Tregs and MDSCs, whereas tumors with low mutational load showed opposite enrichments and depletions (Figure 2b; see also Figure S2 corrections for tumor purity). However, within each mutational load group, the composition of the TILs was divergent for specific cancers as shown in Figure 2c for selected cell types. The tissue context was also observable using correspondence analysis for all immune subpopulations and all cancers (Supplementary Figure S2). Similar trends were observed also within tumor types (Supplementary Figure S3). Additionally, the ratio of effector cells to suppressive cells was also variable. This might explain why some patients with high mutational load are not responsive to therapy with checkpoint blockers and also why some patients with low mutational load are responders.

**Figure 2.**
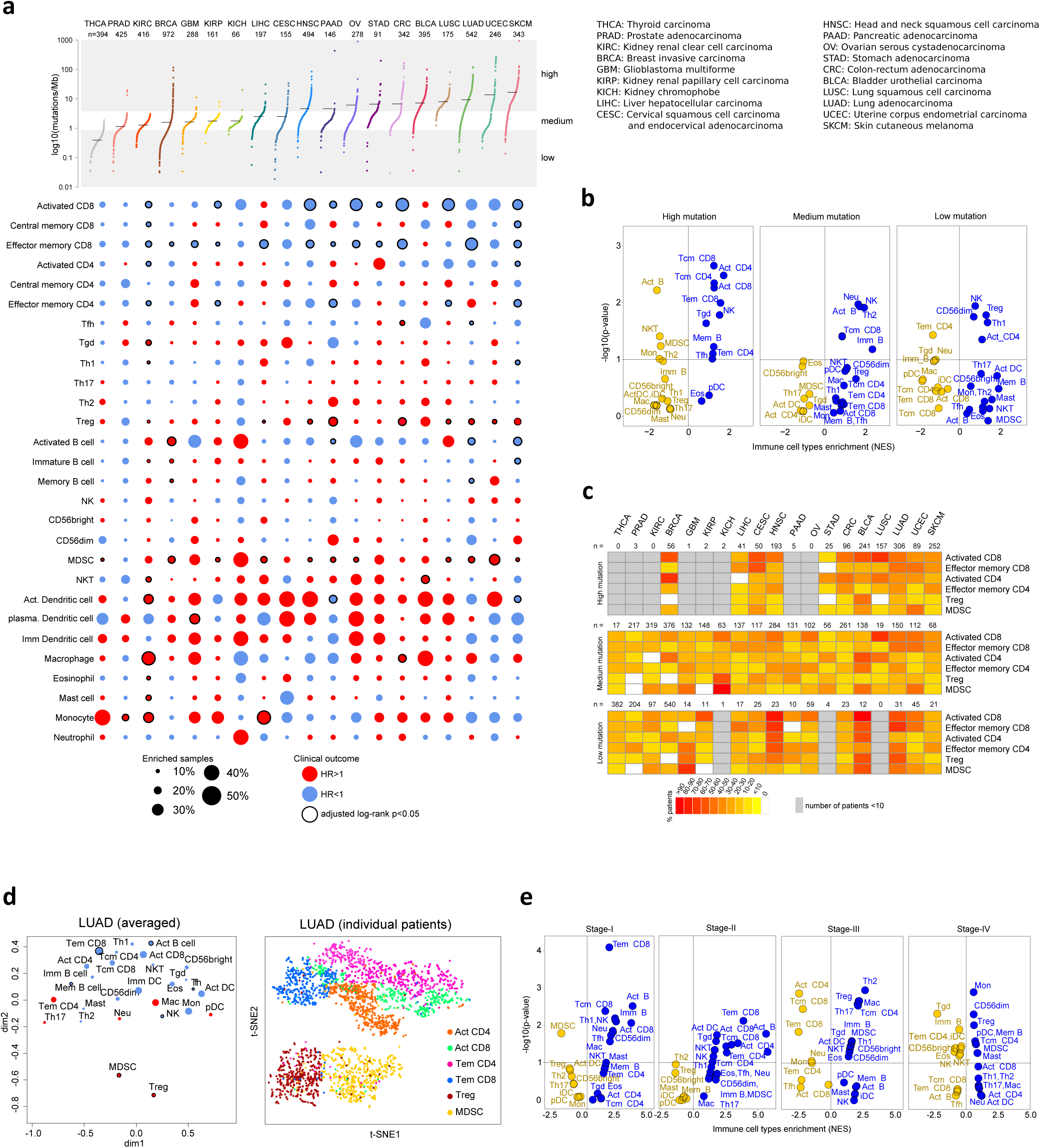
Cellular characterization of immune infiltrates in solid cancers. (**a**) Immune subpopulations across 19 solid cancers. Cancers are sorted according to the mutational load and the immune cell subpopulations alphabetically for adaptive and innate immunity. Lower panel shows results from ssGSEA as bubble plot, where the size of the circles gives the percentage of patients with NES>0 and q-value (FDR)<0.1 and the color indicate the good (blue; HR<1) or bad (red; HR>1) outcome (overall survival). The border indicate adjusted log-rank p-value<0.05. (**b**) Volcano plots for the enrichment (blue) and depletion (yellow) of immune cell types across cancers for tumors with high, intermediate, and low mutational load calculated based on the NES score from the GSEA. (**c**) Fraction of samples in which selected immune subpopulations were enriched in all cancers with high, intermediate, and low mutational load. (**d**) Visualization of the immune infiltrates (averaged normalized enrichment score (NES)) in lung adenocarcinoma (LUAD) for all patients using two dimensional coordinates from multidimensional scaling (MDS) (left panel) and for individual patients and selected cell types based on two dimensional coordinates from t-distributed stochastic neighbor embedding (t-SNE) (right panel). (**e**) Volcano plots for the enrichment (blue) and depletion (yellow) of immune cell types across cancers for tumor stage I to IV calculated based on the NES score from the GSEA.

We then analyzed individual cancer entities and the TILs using dimensional reduction technique and frequently observed separation of the subpopulation related to immune suppression (MDSCs, Tregs) from the subpopulations related to the effector function (activated T cells, Tcm, Tem CD4^+^ and CD8^+^ cells). An example of this is shown in Figure 2d for lung adenoma (LUAD) for all cell types in the LUAD cohort as well as for individual tumors for selected subpopulations. Other examples include ovarian cancer (OV), lung squamous cell carcinoma (LUSC), pancreatic cancer (PAAD), and melanoma (PAAD) (Supplementary Figure S4).

The progression of the tumor across cancers was also characterized by distinct immune cell patterns. For example, Tem CD8^+^ cells were enriched in stage I and stage II tumors and depleted in stage III and stage IV tumors (Figure 2e). In contrast, Tregs and MDSCs were depleted in early stage tumors and enriched in late stage tumors. In general, the enrichment of the TIL subpopulations related to adaptive immunity was decreasing from stage I to stage IV whereas enrichment of TILs related to innate immunity was increasing from stage I to stage IV. This was also evident at the level of individual markers (data not shown). Hence, the cellular composition of the immune infiltrates across solid cancers during progression is shifting towards immunosupressive phenotype.

In summary, these analyses showed that both, the genomic profiles and the specific tissue context contribute to the cellular composition of the immune infiltrates. Furthermore, the results support the notion of the evolving nature of the immune landscape during tumor progression.

### CGAs are associated with infiltration of CD4^+^ and CD8+ T cells

Tumor antigens that have the potential to elicit immune responses that are strictly tumor specific are CGAs and neoantigens (Coulie et al., 2014). CGAs are proteins that are normally expressed by germline cells, but have aberrant expression in tumor cells, whereas neoantigens result from a mutation or rearrangement of a gene-coding sequence. For virus-associated tumors, epitopes derived from viral open reading frames also contribute to the pool of neoantigens (Schumacher and Schreiber, 2015). We therefore characterized these two antigens classes in 20 solid cancers.

The expression of all CGAs (Supplementary Table S2) in all cancers can be retrieved from the website. Since many of these CGAs have low tumoral specificity, we selected 60 CGAs that were previously reported to be transcriptionally silent in normal, non-germline tissues (Rooney et al., 2015) and analyzed the expression and the putative antigens that might be targeted by T cells. In a previous study there was no clear positive association between cytolytic activity and the number of expressed CGAs (Rooney et al., 2015). Here we identified a number of CGAs that were significantly correlated with activated, Tem, and/or Tcm CD8^+^/CD4^+^ cells for each solid cancer (Figure 3a). The number of CGAs significantly associated with T cells ranged between 3 in stomach adenocarcinoma (STAD) and 47 in liver hepatocellular carcinoma (LIHC) (Supplementary Figure S5). The most commons CGA was CAGE1, which was associated with 16 out of the 19 cancers (Figure 3a).

**Figure 3.**
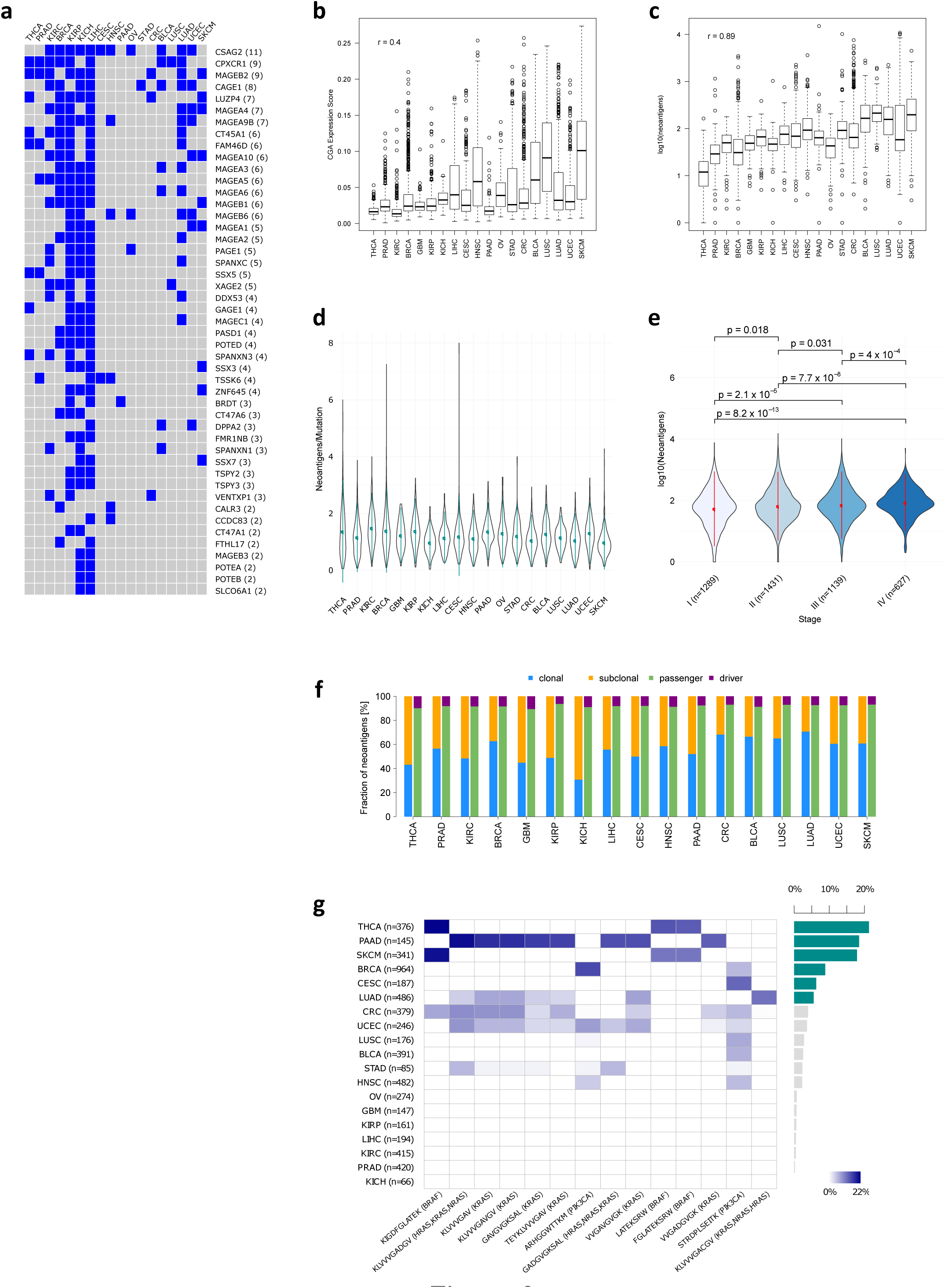
Antigenomes in solid cancers. (**a**) Cancer-germline antigens (CGA) that are associated with CD8^+^ and/or CD4^+^ T cells. Blue squares show significant association with the corresponding cancer type (Spearman rank correlation>0.3; adjusted p-value<0.1). CGAs are sorted according to the number of cancers with significant associations. (**b**) CGA expression score across cancers. Cancers are sorted according to the mutational load (**c**) Neoantigen load across cancers. (**d**) Neoantigen frequencies for solid cancers. (**e**) Neoantigen load for different tumor stages (Kruskal Wallis test followed by two-sided Dunn’s pairwise post hoc tests on rank sums with Benjamini-Hochberg adjustment of p-values). (**f**) Fractions of neoantigens and their origin. (**g**) Shared neoantigens in solid tumors. Shown are only neoantigen shared in at least 5% of the tumors.

As these CGAs are candidates for vaccination we also predicted the peptides from these CGAs that bind to HLA molecules, leading to 5775 unique peptides (Supplementary File). We then estimated the antigen load by calculating a CGA expression score (Computational methods). Interestingly, there was positive correlation of the CGA score with mutational load across cancers (r=0.4, p<2.2e−16) (Figure 3b).

These results suggest a tissue-dependent role of CGAs in spontaneous immunity and reinvigorate the use of CGA-based therapeutic vaccines, perhaps in combination with epigenetic drugs to increase the expression of selected CGAs.

### Neoantigen landscape is diverse and sparse

As neoantigens are attractive candidates for developing therapeutic cancer vaccines, we analyzed the neoantigens landscape in order to identify potential candidates for off-the-shelf vaccination. The neoantigen landscape in solid tumors was composed of 933,954 expressed neoantigens (911,548 unique) originating from 893,960 somatic point mutations. As expected, the number of neoantigens correlated with the mutational load (Figure 3c). The median neoantigen frequency, i.e. the number of neoantigens per mutation varied across cancers and ranged from 0.93 (skin cutaneous melanoma (SKCM) to 1.43 (kidney renal cell carcinoma (KIRC)) (Figure 3d). It should be noted that these differences across cancers might be attributed to the differences in mutational processes, which are more likely to lead to nonsynonymous mutations (e.g. C>A transversions compared to C>T transversions (Lawrence et al., 2013)). The associations of the neoantigen load as well as of the CGA load and selected genes and immune cell subpopulations were highly variable between different cancer entities (Supplementary Figure S6), suggesting that the immune response is likely governed by combined effects of both antigen classes.

Somewhat unexpectedly, there was only a slight increase of the neoantigen load from stage I to stage IV across all cancers (all adjusted pairwise p-values<0.032; two-sided Dunn’s post hoc tests on ranked sums; Figure 3e). Furthermore, as the cellular composition of the immune infiltrates during progression is shifting towards immunosuppressive phenotype this raises an issue of whether immunotherapies that depend on the adaptive immune response can be effective in later stage.

The fraction of neoantigens derived from driver genes for all solid tumors was 7.6% (ranging between 7.0% for LUSC and 10.6% for glioblastoma (GBM)) (Figure 3f). Hence, the bulk of neoantigens had its origin in passenger genes. On average, 56% of the neoantigens were of clonal origin with varying proportions of clonal to subclonal fractions across cancers (Figure 3f). The fractions of neoantigens with clonal origin ranged from 31% (KICH) to 71% (LUAD) (Supplementary Table S3).

The neoantigens were infrequently shared between patients (Figure 3g; see also Supplementary Figure S7 for HLA-stratified shared neoantigens). From the total of 911,548 unique neoantigens only 24 were shared in at least 5% of patients in one or more cancer types. These shared neoantigens represent identical peptides originating from one or more genes. As expected, the most frequent neoantigens were induced by mutations in driver genes like BRAF, RAS, and PIK3CA. Among these, only two peptides were shared in more than 15% of patients in one cancer type: KIGDFGLAT**<u>E</u>**K was shared in THCA and SKCM, and KLVVVGA**<u>D</u>**GV was shared in PAAD. The neoantigen KIGDFGLAT**<u>E</u>**K originates from BRAF^V600E^ mutation, which is present in a large fraction of THCA (Cancer Genome Atlas Research, 2014) and SKCM (Cancer Genome Atlas, 2015) tumors. The neoantigen KLVVVGA**<u>D</u>**GV originates from the p.G12D mutation of KRAS, which is shared across a large fraction of PAAD patients (Witkiewicz et al., 2015).

Thus, the antigenome landscape was diverse between and within cancers, and with respect to the neoantigens highly sparse. Furthermore, the results suggest that T cell responses are not only directed against neoantigens, but also against CGAs

### Genotypes of the tumors determine immunophenotypes and tumor escape mechanisms

The immunogenomic analyses of the CRC data in our previous study revealed that the immunophenotypes in the hypermutated compared to the non-hypermutated tumors were characterized by increased enrichment of effector T cells (Angelova et al., 2015), likely to be a consequence of the higher antigen burden. We asked the question how are the immunophenotypes related to other genomic features describing the complexity of the tumor genome, like tumor heterogeneity (high vs. low) and antigenicity (high vs. low).

Tumors that were more heterogeneous were enriched with activated T cells and Tem cells, and depleted of immunosuppressive cells, despite slightly lower mutational load (Figure 4a and 4b). Tumors with higher neoantigen frequencies showed similar cellular patterns of immune infiltrates (Figure 4c and 4d). We then asked if there is a genotype-immunophenotype relationship also for gene-specific view of the genomic landscape. We selected two cancers with the lowest and highest mutational load, i.e. THCA and SKCM, and analyzed the immunophenotypes for their distinct genotypes: BRAF and RAS subtypes for THCA (Cancer Genome Atlas Research, 2014), and BRAF, RAS, NF1, and triple negative wild-type for SKCM (Cancer Genome Atlas, 2015). Additionally, we examined the expression of three classes of molecules that are involved in tumor escape mechanisms: MHC molecules (class I, class II and non-classical) which may be downregulated to avoid recognition by T cells, immunostimulators (e.g. OX40), which may be down-regulated to avoid immune destruction, and immunoinhibitory genes (e.g. CTLA-4) which may be up-regulated to enable tumor escape.

**Figure 4.**
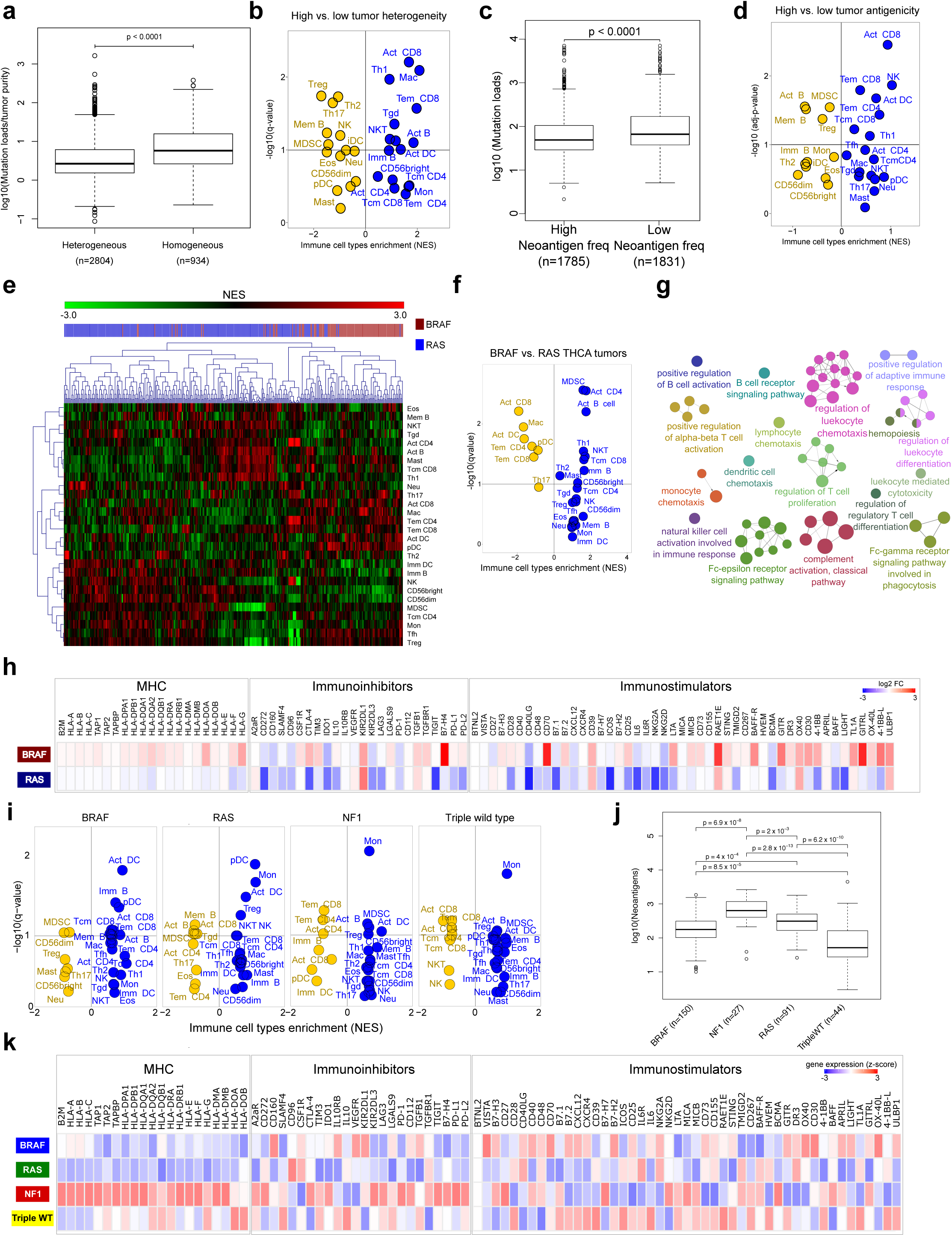
Genotypes and immunophenotypes in solid cancers. (**a**) Mutational load and tumor heterogeneity (two-sided Wilcoxon rank sum test) (**b**) Immune infiltrates in tumors. Shown is a volcano plot for tumors with high and low heterogeneity calculated based on the NES score from the GSEA. (**c**) Mutational load and neontigen frequency (two-sided Wilcoxon rank sum test). (**d**) Immune infiltrates in tumors. Shown is a volcano plot for tumors with high and low antigenicity calculated based on the NES score from the GSEA. (**e**) Hierarchical clustering of immune cell composition for BRAF and RAS mutated THCA tumors. (**f**) Volcano plot for BRAF and RAS mutated TCHA tumors calculated based on the NES score from the GSEA. (**g**) Gene ontology (GO) analysis of the differentially expressed genes for BRAF and RAS mutated TCHA tumors using ClueGO (Bindea et al., 2009). (**h**) Expression of MHC and immunomodulatory molecules in BRAF and RAS mutated TCHA tumors. Expression values were compared to normal tissue (log_2_-fold changes are color coded according to the legend). (**i**) Volcano plots for SKCM genotypes calculated based on the NES score from the GSEA. (**j**) Mutational load for SKCM genotypes (Kruskal Wallis test followed by two-sided Dunn’s pairwise post hoc tests on rank sums with Benjamini-Hochberg adjustment of p-values). (**k**) Expression of MHC and immunomodulatory molecules for SKCM genotypes. Expression values are represented by z-score calculated across all SKCM tumors and color coded according to the legend

The different THCA genotypes were associated with specific immunophenotypes (Figure 4e) and showed distinct cellular patterns of immune infiltrates (Figure 4f) in spite of comparable neoantigen burden (p>0.05, two-sided Wilcoxon rank sum test). Analysis of the differentially expressed genes with respect to the immune-related Gene Ontology (GO) terms highlighted the pathways, which might explain the different effects on the immune system including chemotaxis, T cell differentiation, T cell proliferation, T cell activation, or B cell receptor signaling pathway (Figure 4g). The expression levels of MHC molecules, immunostimulatory and immunoinhibitory molecules were also associated with the genotypes (Figure 4h). These results suggest that BRAF mutated THCA tumors employ different tumor escape mechanisms compared to RAS mutated tumors in THCA: BRAF tumors were infiltrated with immunosuppressive cells, whereas RAS tumors downregulated MHC molecules as well as immunomudulatory molecules.

Similarly, the SKCM genotypes were also associated with distinct immunophenotypes (Figure 4i), with BRAF tumors enriched with effector T cells, whereas other genotypes were enriched with immunosuppressive cells. It is noteworthy that for the SKCM cohort the neoantigen burden differed (all adjusted pairwise p-values ≤ 0.002; two-sided Dunn’s post hoc tests on ranked sums; Figure 4j), and therefore the relative contribution of the mutational load can have an additional impact. The four SKCM genotypes were also associated with varying expression levels of MHC and immunomodulatory molecules (Figure 4k). Analysis of the differentially expressed genes with respect to the immune-related GO terms is shown in the Supplementary Material (Supplementary Figure S8).

The results of this study suggest that the genotypes of the tumor determine the immunophenotypes and the tumor escape mechanisms. This was evident at the high-level view of the genomic landscape (e.g. mutational load, tumor heterogenity) as well as at the gene-specific view (e.g. BRAF or RAS mutated tumors).

### Machine learning identifies major determinants of tumor immunogenicity in solid cancers

The results of this work showed not only highly heterogeneous TILs but also varying ratios of different T cell subsets including suppressive ones. These observations raise the question of the underlying molecular mechanisms that explain the differences in immunogenicity of the tumors. The question can be reduced to the notion of sources of immunogenic differences, which can be divided into two categories: tumor intrinsic factors and tumor extrinsic factors. Tumor intrinsic factors include the mutational load, the neoantigen load, the neoantigen frequency, the expression of immunoinhibitors and immunostimulators (e.g. PD-L1 (Coulie et al., 2014)), and HLA class I molecule alterations (Garrido et al., 2010). Tumor extrinsic factors include chemokines which regulate T cell trafficking (Gajewski et al., 2006), infiltration of effector TILs and immunosuppressive TILs, and soluble immunomodulatory factors (cytokines) (Gajewski et al., 2006).

In a previous study an association of the cytolytic activity and the neoantigen load was reported (Rooney et al., 2015). Here, we asked the question what are the determinants of the tumor immunogenicity and considered all tumor intrinsic and tumor extrinsic factors mentioned above. The number of single parameters that determine the immunogenicity of the tumors and the various combinations thereof render it difficult to identify a set of representative parameters even with large cohorts like the TCGA ones. We therefore employed machine learning techniques and exploited the large-scale genomics data. We reasoned that the immunogenicity of the tumor can be represented by the cytolytic activity (estimated using the expression of granzyme A (*GZMA*) and perforin (*PRF1*) according to (Rooney et al., 2015)), as this is the ultimate effector mechanism in the cancer immunity cycle (Chen and Mellman, 2013). For each cancer type we used a random forest classification approach, which is based on a multitude of decision trees, including 127 parameters (Supplementary Table S4) to separate tumors with high cytolytic activity from tumors with low cytolytic activity. For individual cancer types the most predictive features were identified using the mean decrease of accuracy over all cross validated predictions (Figure 5a). For each of the studied cancers the analysis revealed only immune-related factors which we classified into four categories: 1) infiltration of activated CD8^+^/CD4^+^ T cells and Tem CD8^+^/CD4^+^ cells, 2) infiltration of immunosuppressive cells (Tregs and MDSCs), 3) expression of MHC class I, class II, and non-classical molecules, and 4) expression of certain co-inhibitory and co-stimulatory molecules (Figure 5a).

**Figure 5.**
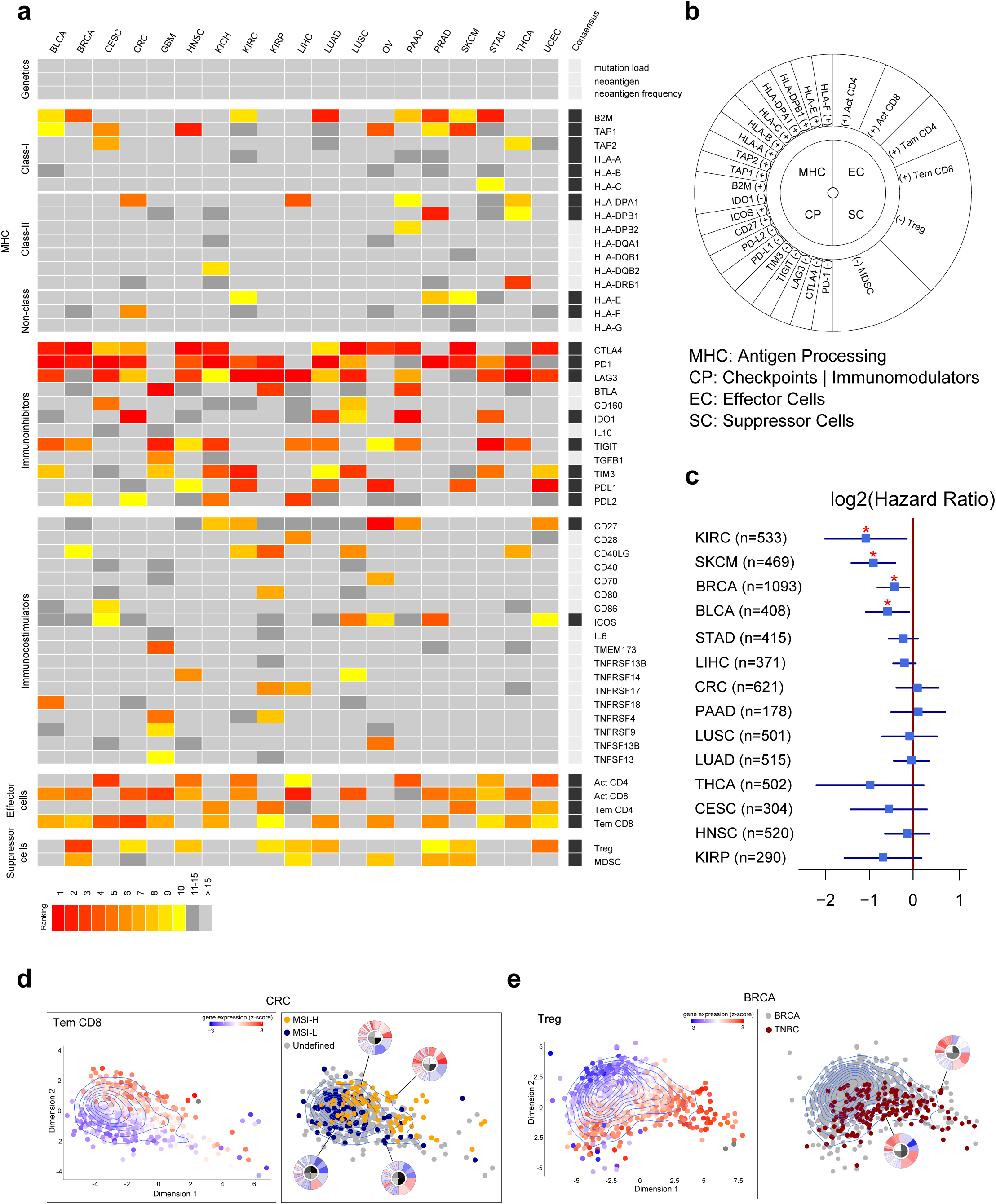
Determinants of immunogenicity in solid cancers. (**a**) Major parameters determining immunogenicity in solid cancers revealed using random forest approach. (**b**) Immunophenogram for the visualization of the parameters determining immunogenicity. (**c**) Results of multivariate survival analyses using the immunophenoscore for all solid cancers. Forest plots showing log2 hazard ratio (95% confidence interval), * indicates adjusted p<0.05 (**d**) Visualization based on two dimensional coordinates from multidimensional scaling (MDS) of expression profiles from the genes and the gene sets as used in the immunophenogramm in colorectal cancer (CRC). Mean expression (z-score) of immune genes of Tem CD8^+^ T cells as well as MSI and MSS samples for CRC are indicated by colors according to the legend and the mean expression (z-scores). (**e**) Visualization based on two dimensional coordinates from MDS in breast cancer (BRCA). Tregs and triple negative breast cancer (TNBC) samples for BRCA are indicated by colors according to the legend. Representative immunophenograms for selected patients are shown.

To visualize the information about the immunophenotypes of the tumors based on the identified major determinants we constructed an immunophenogram that includes these four categories (Figure 5b). We then calculated an aggregated score – immunophenoscore – based on the expression of the representative genes or gene sets comprising four categories: MHC molecules, immunomodulators, effector cells (activated CD8^+^ T cells and CD4^+^ T cells, Tem CD8^+^ and Tem CD4 ^+^ cells), and suppressor cells (Tregs and MDSC) (see Computational Methods). Multivariate analysis showed that the immunophenoscore was associated with survival in 12 solid cancers, of which four were significant: KIRC, SKCM, BRCA, and bladder cancer (BLCA) (Figure 5c).

The immunophenogram enables the visualization of the immunophenotypes of the tumor also at the levels of individual tumors as seen for CRC (Figure 4d) and BRCA (Figure 4e). As can be seen, subgroups of tumors like microsatellite-instable (MSI) and triple-negative breast cancer (TNBC) are grouped with respect to distinct TILs like Tem CD8^+^ cells and Tregs, respectively. We also implemented an interactive version of the tool (see <u>http://tcia.at</u>), which enables visualization and scoring of tumor samples using expression data for the markers we identified using the random forest approach.

Hence, using a data-driven approach, a large number of tumors (>8000), and genomic, transcriptomic, and immunological features, we were able to identify the major determinants of tumor immunogenicity. Based on these results we propose a visualization method – the immunophenogram, and a scoring scheme – the immunophenoscore, for solid tumors.

### Immunophenoscore predicts response to immunotherapy with CTLA-4 and PD-1 blockers

As only a minority of the patients is responsive to checkpoint blockers, the identification of predictive markers and the mechanisms of resistance to immunotherapy is a subject of intense research. Although several markers have been proposed including TILs, PD-1 or PD-L1 expression, mutational load (Rizvi et al., 2015) or clonal neoantigens (McGranahan et al., 2016), none have yet been fully validated (Spencer et al., 2016).

We reasoned that the determinants of immunogenicity identified using the random forest approach might have also a predictive value and analyzed two genomic and transcriptomic data sets from patients with melanoma treated with anti-CTLA-4 (Van Allen et al., 2015) and anti-PD-1 antibodies (Hugo et al., 2016). Using RNA-sequencing data and GSEA we reconstructed the TIL landscape and scored the patients using the immunophenoscore. The immunophenograms of the individual patients treated with anti-CTLA-4 antibodies are shown in Figure 6a. Tumors of the responders were enriched with cytotoxic cells (CD8^+^ T cells, Tγδ, NK cells) and depleted of MDSCs and Tregs (Figure 6b). More importantly, the immunophenogram and the score derived from the analyses enabled stratification of patients to responders and non-responders (Figure 6c) with a superior predictive power compared to the expression of checkpoint molecules as can be seen in the receiver operating characteristics (ROC) curve (Figure 6d).

**Figure 6.**
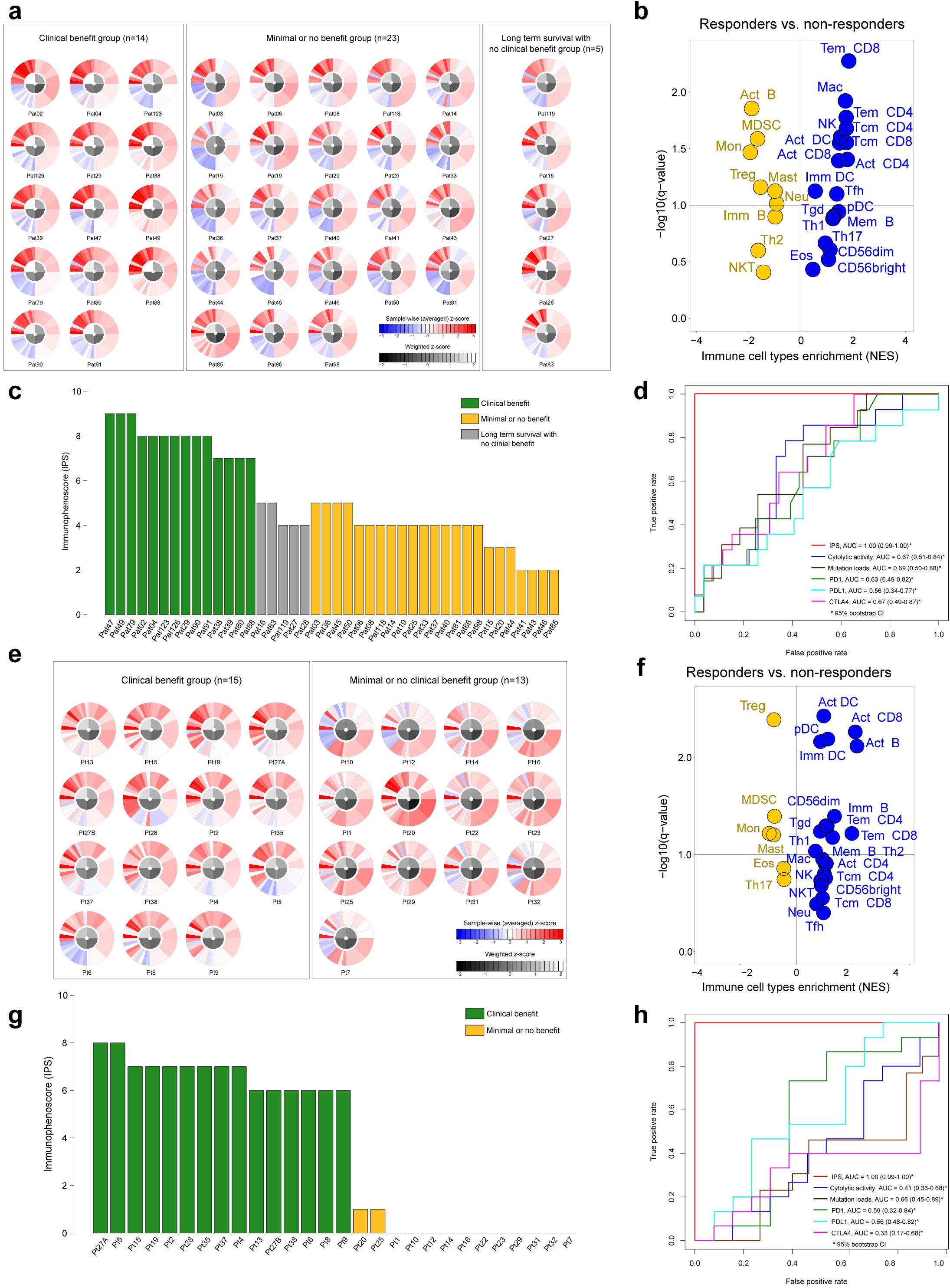
Immunophenoscores (IPS) and response to checkpoint blockade. (**a**) IPS and response to blockade with anti-CTLA-4 antibody (Data from van Allen et al.(Van Allen et al., 2015)). Shown are immunophenograms for individual patients. HLAs were highly upregulated compared to the mean expression within the tumor (z>0; indicated red within the upper left quadrant termed MHC), whereas many checkpoints were downregulated compared to the mean expression with that tumor (z<0; indicated blue in the lower left quadrant termed CP) in many patients of this study. (**b**) Volcano plot for the enrichment and depletion of immune subsets in the tumor calculated based on the NES score from the GSEA. (**c**) IPSs for the cohort. (**d**) Receiver operating characteristics for IPS, and for the cytolitic activity (mean expression of GZMA and PRF1 as suggested by (Rooney et al., 2015)), mutational load, and for the expression of PD-1, PD-L1, and CTLA4. AUCs with 95% bootstrap confidence interval are provided. (**e)** IPS and response to blockade with anti-PD-1 antibody (Data from Hugo et al. (Hugo et al., 2016)). Shown are immunophenograms for individual patients. (**f**) Volcano plot for the enrichment and depletion of immune subsets in the tumor calculated based on the NES score from the GSEA. (**g**) IPSs for the cohort. (**h**) Receiver operating characteristics for IPS, and for the cytolitic activity (mean expression of GZMA and PRF1 as suggested by (Rooney et al., 2015)), mutational load, and for the expression of PD-1, PD-L1, and CTLA4. AUCs with 95% bootstrap confidence interval are provided.

Similar observation was made also for the patients treated with anti-PD-1 antibody. Visualization of the determinants of the immunogenicity with the immunophenogram for responders and nonresponders showed distinct expression patterns in the two groups (Figure 6e). Again, tumors of the responders were enriched with cytotoxic cells (CD8^+^ T cells, Tγδ, NK cells) and depleted of MDSCs and Tregs, as evident in the volcano plot (Figure 6f). Finally, the immunophenoscore (Figure 6g), and the ROC curve (Figure 6h) showed the predictive value also for patients treated with anti-PD-1 antibodies.

Hence, the immunophenoscore developed from a panel of immune-related genes belonging to the four classes: effector cells, immunosuppressive cells, MHC molecules, and selected immunomodulators has predictive value in melanoma patients treated with the CTLA-4 and PD-1 blockers.

## DISCUSSION

We used an analytical strategy and for the first time provide comprehensive view of 28 TIL subpopulations including effector and memory T cells, and immunosuppressive cells (Tregs, MDSCs). We also developed an interactive big data resource that includes cellular compositions of TILs, neoantigens, CGAs, and tumor heterogeneity, and enables other researchers to generate testable hypotheses. Our approach to deeply mine cancer genomic datasets revealed a number of novel associations suggesting important biological conclusions with implications for cancer immunotherapy. Moreover, using machine learning approach we identified major determinants of tumor immunogenicity and propose an immunophenoscore, which is not only prognostic like the immunoscore for CRC (Mlecnik et al., 2011), but has also superior predictive power for identifying responders for treatment with CTLA-4 and PD-1 antibodies.

First, our intriguing observation of the association of the genotypes and the immunophenotypes and the distinct tumor escape mechanisms determined by the genotypes implicates that the interaction with the immune system is predetermined by the genetic basis of the tumors. This genotype-immunophenotype relationship was evident at the high-level view of the genomic landscape, i.e. heterogeneous vs. homogenous, and antigenic vs. less antigenic tumors. Strikingly, this association was observed also at the gene-specific view, i.e. at the level of mutational origin of the tumors like BRAF or RAS, as well as for tumors at both ends of the mutational load: THCA and SKCM. This raises the question of the underlying molecular mechanisms. Currently we can only speculate on the possible mechanisms, which can include modifications of different signaling pathways due to mutational changes. For example, evidence from clinical and experimental studies has demonstrated that signaling by BRAF and RAS oncogenes can present similarities but also differential effects (Oikonomou et al., 2014). The analysis of the genes differentially expressed between the two genotypes in THCA indicates several processes that might be involved in the sculpting of the immune landscape (Figure 4g).

Second, charting of the antigenome showed that both, the CGAs and the neoantigens are associated with effector and memory T cells, suggesting that T cell responses are directed against both antigen classes. Moreover, the antigen load of the CGAs and neoantigens was highly variable across and within cancers. While neoantigens are thought to be of particular relevance to tumor control (as these antigens are not affected by central T cell tolerance) (Schumacher and Schreiber, 2015), the contribution of the CGAs is still unclear. In a previous study the role of the CGAs in anti-tumor immunity was not uncovered (Rooney et al., 2015), likely due to the methods used (e.g. expression signatures of only two genes). In contrast, our comprehensive and high-resolution approach provides evidence for T cell reactivity also against CGAs. The quality of the T cell responses directed against neoantigens and CGAs has not been compared so far and future experimental studies are needed to identify the contribution of each class of antigens. We envision that in case of malignancies with a low mutational load that are less likely to produce immunogenic neoantigens, CGAs might be therapeutically exploited by combining DNA hypomethylation agents to induce expression of CGAs and checkpoint blockers to target tumor cells. This type of combination strategy was recently evaluated in a mouse model (Covre et al., 2015).

Beyond these biological insights, the results from this study have also important implications for cancer immunotherapy with checkpoint blockers as monotherapy, as combination therapy with targeted agents, and for therapeutic vaccination. Most importantly, we propose an immunophenoscore, i.e. a panel of immune genes for classification of patients likely to respond to therapy with antibodies targeting CTLA-4 and PD-1 with superior performance. The immunophenoscore we developed was derived in an unbiased manner using the TCGA data and machine learning, but it reflects current understanding of the categories of genes that determine immunogenicity of the tumors: effector cells, immunosuppressive cells, MHC molecules, and immunomodulators. The immunophenoscore is similar to the conceptual immunogram that was recently proposed to represent the status of the immune system (Blank et al., 2016). Another advantage of the immunophenoscore is that it represents a standardized value since z-scores are used and is therefore more robust compared to the use of expression values. However, since as of today only limited data is available, additional studies are required to validate the immunophenoscore. Notably, the method can be further improved by optimizing the immunophenoscore for specific cancers. Finally, for routine applications other techniques for gene expression profiling like microarrays and qPCR can be used instead of RNA sequencing.

As only a minority of the patients is responsive to checkpoint blockers, there are major efforts to develop therapeutic strategies to overcome resistance by using combination of checkpoint blockers and targeted agents. Hence, precision immune-oncology, i.e. the amalgamation of immunotherapy and precision oncology appears as a promising approach to treat patients. Supported by insights from genomic profiling studies and the availability of a large number of drugs and molecular targets, the precision oncology hypothesis was postulated, i.e. that cancer treatment decisions could be guided by molecular profiling irrespective of the tissue of origin. Since the efficacy of checkpoint blockers was shown in several cancers, one could adopt this tissue-agnostic approach also for precision immune-oncology. However, the results of our analyses show that the tissue context is an important determinant of the cellular composition of the immune infiltrates, and hence, determines the antitumor response. Consequently, stratification of patients in a precision immune-oncology framework requires genomics-based information and characterization of the immune infiltrates.

With respect to neoantigen-based therapeutic vaccination, the sparsity of the neoantigen space advocates against the development of off-the-shelf vaccines. A notable exception represents vaccine for THCA, SKCM or PAAD, which could target 20% of the patients. Thus, personalized cancer vaccination strategy is required in which whole-exome NGS is performed to identify somatic mutations, followed by bioinformatics analyses to identify neoantigens, and synthesis of peptide- or DNA/RNA-based vaccines. Viability of such personalized cancer vaccination strategy was recently demonstrated in clinical studies in melanoma patients (Robbins et al., 2013; van Rooij et al., 2013).

Finally, with the large number of ongoing studies with checkpoint blockers either as monotherapy or combination therapy, we expect that the immunogenomic data amount will continuously increase. Thus, we strongly believe that the TCIA in its current form and future incorporation of additional datasets represents an important contribution to the field and will enhance the identification of novel mechanistic insights of the complex tumor-immune cell interactions.

## COMPUTATIONAL METHODS

### Identification of immune-related genes

For the identification of immune-related genes we used Affymetrix HG-U133A microarray data from a number of different studies (Supplementary Table S5) according to our recently developed approach for TILs in CRC (Angelova et al., 2015), and included expression profiles from normal tissues (Petryszak et al., 2016) and from relevant cancer cell lines (Barretina et al., 2012). The so defined immune-related gene expression signatures comprised 1625 genes. For the identification of subsets of genes representative for specific immune cell types, we selected genes with an average correlation r ≥ 0.4 (p < 0.01) between all specific immune genes in the same cell type. This threshold was chosen to satisfy two goals: selection of genes with relatively high correlation such that their correlation could not be considered a chance event; and selection of a reasonable number of genes (at least 10 genes per subpopulations). Furthermore, using a set of genes instead of individual markers for specific TILs ensures robust estimation and is less susceptible to noise arising from the expression of the genes in tumor or stromal cells. The 782 immune genes 782 are listed in the Supplementary Table S6.

### Genomic and clinical data

Genomic and clinical data for 20 solid tumors from The Cancer Genome Atlas (TCGA) were downloaded via the TCGA data portal or queried via Broad GDAC firehose/firebrowse and include clinical information (n=9151), SNP arrays (n=3377), microarrays (n=550), and RNA-sequencing expression profiles (n=8243), as well as curated MAF files of somatic mutations (Kandoth et al., 2013) (Supplementary Table S1). All curated MAF files were re-annotated with Oncotator (Ramos et al., 2015) in order to have a common annotation resource and file format. Notably, as we used the latest available information, our results differ slightly from previously published ones (Alexandrov et al., 2013) (e.g. due to different numbers of samples we observed different mutational loads for LUAD (n=542 vs. 381) and LUSC (n=178 vs. 176)). Mutational load was also corrected for tumor purity (Supplementary Figure S9). FASTQ files of RNA-sequencing reads (n=8,398) were downloaded from CGHub (<u>https://cghub.ucsc.edu</u>) and analyses were done on the fly. RNA-sequencing expression levels, available as Transcripts-Per-Millions (TPM) were transformed to log_2_(TPM+1). For RNA-sequencing expression data, we considered data from primary solid tumor and normal solid tissue for all cancer types. In case of subjects with multiple RNA-sequencing samples available, the sample with the highest sequencing depth was considered for GSEA and deconvolution.

### Identification of TILs subpopulations

We used single sample gene set enrichment analysis (ssGSEA) (Barbie et al., 2009) to identify immune cell types that are over-represented in the tumor microenvironment. The expression levels of each gene were z-score normalized across all patients. For each patient (or group of patients) genes were then ranked in descending order according to their z-scores (mean of z-scores). The association was represented by a normalized enrichment score (NES). An immune cell type was considered enriched in a patient or group of patients when FDR (q-value) ≤ 10%. Clustering and visualization was done with the software Genesis (Sturn et al., 2002). Enriched immune cell types (NES>0) are illustrated as bubble plots (similar to (Spinelli et al., 2015)) where the size represents percentage of patients with enriched cell type and color is encoded by hazard ratio from overall survival analysis. The similarity of the enrichment of immune infiltrates (averaged NES) were calculated using multidimensional scaling (MDS). The distribution of selected cell types for individual patients were analyzed with t-distributed stochastic neighbor embedding (t-SNE) (van der Maaten and Hinton, 2008) using the Matlab toolbox t-SNE.

Additionally, a deconvolution approach was applied using the tool CIBERSORT (Newman et al., 2015) using a custom model to modify RNA-sequencing data from TCGA to be used as input of the deconvolution algorithm. To build the model, we considered only tumor samples for which both Affymetrix microarrays and Illumina RNA-sequencing data were available (131 samples for LUSC, 266 for OV and 153 for GBM). We summarized RNA-seq expression as log2(TPM+1) and removed genes with null expression in more than 50% of samples. We modeled the RNA-sequencing-to-microarrays mapping with a gene-specific, cubic smoothing spline with four degrees of freedom and used the model to transform RNA-sequencing data into microarrays-like data. Transformed data were then processed with the R-based version of CIBERSORT, with default parameter settings. The performance of the model was tested on these three cancer types with leave-one-out cross-validation, confirming a high agreement between the cell fractions estimated from the modified RNA-sequencing data and those computed from the corresponding microarray data (0.88 Pearson’s correlation, data not shown).

### Cancer-germline antigens

The list of cancer germline antigens (CGA) was extracted from the Cancer-Testis database (Almeida et al., 2009) (Supplementary Table S2). The heatmap of CGA expression per cancer type was computed on the median log_2_(TPM+1) values per cancer type considering only expressed CGAs (median TPM per cancer type higher than 2). The list of tumor-specific CGAs was adopted from Rooney et al. (Rooney et al., 2015). We computed the CGA Expression Score for each cancer patient, defined as the area under the curve of the fraction of CGA exceeding increasingly higher expression thresholds (computed only on tumor-specific CGAs).

CGA protein sequences were retrieved from Uniprot (<u>http://www.uniprot.org/</u>). All possible 9nt-long peptides spanning CGA proteins were subjected to NetMHCpan 3.0 (Nielsen and Andreatta, 2016) to predict their binding affinity to each of the HLAs estimated for TCGA patients. Only peptides with a predicted binding affinity lower than 500 nM was considered (Supplementary File.)

### Characterization of neoantigens

HLA alleles were called from RNA-sequencing FASTQ files using Optitype (Szolek et al., 2014), selected for its high performance and for its applicability to RNA-sequencing data. For each subject, the HLA alleles estimated from the sample with the highest coverage over the HLA locus were considered. To estimate the mutated proteins, we focused on non-synonymous missense mutations, and selected mutations associated to Uniprot protein identifiers. The protein sequence retrieved from Uniprot was changed according to the non-synonymous, missense mutations reported in the MAF file, and truncated in case of stop codons. We removed candidate proteins affected by annotation inconsistencies between protein identifiers and predicted effect. Peptides of 8-11 amino acids in length, covering the mutated region of the protein, were analyzed with NetMHCpan (Nielsen et al., 2007) (Version 2.8) to estimate their binding affinity to the HLA alleles. Self-antigens mapping to human Uniprot proteins were identified with BLAST and filtered out. Amongst the candidate antigenic peptides, we selected strong binders with binding affinity < 500 nM as in (Rooney et al., 2015), and considered peptides arising from expressed genes. We identified expressed genes as those having median TPM greater than 2 in a given cancer type, as previously shown (Wagner et al., 2013). By plotting the distribution of TPM expression of all genes in TCGA tumor samples, we verified that this threshold provides a good separation between two population of genes: one genes with null or low expression, possibly corrupted by technical noise, and one mid-to-high expression genes (see Supplementary Figure S10).

### Estimation of tumor heterogeneity and clonality of mutations

The ABSOLUTE algorithm (Carter et al., 2012) was used to integrate the copy number data together with the somatic mutations in order to estimate the purity and ploidy, and measure the fraction of cancer cells per mutation (CCF). The SNP data were downloaded from the TCGA portal and analyzed with HAPSEG (Carter et al., 2011). The tumor heterogeneity was estimated as the area under the curve (AUC) of the cumulative density function from all cancer cell fractions per tumor. Tumors were considered homogenous within the lower quartile of the AUC distribution whereas all other tumors were considered heterogeneous. A mutation was classified as clonal if the CCF was > 0.95 with probability > 0.5, and subclonal otherwise (Landau et al., 2013). Since at the time of our data freeze, only hg18 was available for OV and the SNP array data for STAD were not processed, these cancers were excluded from the analysis of the clonal/subclonal origin of neoantigens.

### Identification of determinants of tumor immunogenicity, immunophenogram, and immunophenosocore

For each patient the cytolytic activity was calculated as the mean of the *GZMA* and *PRF1* expression levels (log2 (TPM+1)) as previously defined (Rooney et al., 2015). For each cancer type, patients were divided into two groups based on median cytolytic activity. A random forest classifier (Breiman, 2001) separating the group of patients with TILs exhibiting higher cytolytic activity from the group of patients with TILs exhibiting lower cytolytic activity was trained using the R package *randomForest* with 10,000 trees and included mutational load per megabase, number of neoantigens, fraction of neoantigens per mutations, expression of MHC related molecules, expression of immunomodulatory factors, and mean expression of the respective immune genes for each of the 28 immune celltypes as independent variables. Analysis of the association between TILs and MHC molecules showed varying associations (Supplementary Figure S11). The mean decrease of accuracy over all out-of-bag cross validated predictions was used to rank predictors.

The immunophenogram was constructed similar to the recently proposed metabologram (Hakimi et al., 2016). From the results of the random forest approach a list of consensus determinants (Figure 5a) that includes 20 single factors (MHC molecules, immunoinhibitors, and immunostimulators) and six cell types (activated CD4^+^ T cells, activated CD8 ^+^ T cells, effector memory CD4^+^ T cells, and effector memory CD8^+^ T cells, Tregs, and MDSCs) were selected for further analyses. For each determinant a sample-wise z-score from gene expression data was calculated. For the 6 cell types an average z-score from the corresponding metagenes was calculated. The determinants were then divided into four categories: effector cells (activated CD4^+^ T cells, activated CD8^+^ T cells, effector memory CD4^+^ T cells, and effector memory CD8^+^ T cells), suppressive cells (Tregs, MDSCs), MHC related molecules, and checkpoints/immunomodulators, and color-coded in the outer part of the wheel (red: positive z-score, blue: negative z-score). The z-scores of the determinants included in the particular category were positively weighted with the weight one (each MHC molecule, ICOS, CD27, activated CD4^+^ T cells, activated CD8^+^ T cells, effector memory CD4^+^ T cells, and effector memory CD8^+^ T cells) and negatively weighted with the weight one (PD-1, CTLA4, LAG3. TIGIT, TIM3, PD-L1, PD-L2, Tregs, MDSCs). The weighted averaged z-score score was then calculated by averaging the z-scores within the respective category leading to four values, which were subsequently gray scaled. The immunophenscore (IPS) was calculated on an arbitrary 0-10 scale based on the sum of the weighted averaged z-score of the four categories, whereby the sum of the z-scores≥3 were designated as IPS10 and the sum of the z-scores≤0 are designated as IPS0.

To determine the predictive power patients were stratified into responders and non-responders and a ROC validation including the IPS, cytolytic activity, CTLA-4 expression, PD1 expression, PD-L1 expression as independent variable was performed. As predictive value the area under the curve (AUC) from the ROC analyses was used (R package *ROCR)*. For each of the analyzed patients the corresponding immunophenogram is provided in the TCIA. A tool for construction of the immunophenogram on other expression data was implemented in the TCIA and a respective R-code is available at github (<u>https://github.com/MayerC-imed/Immunophenogram</u>).

### Statistical analyses

Sample sizes from available TCGA data were considered adequate as sufficient power using equivalent tests was observed in a previous study (Angelova et al., 2015). To test for differential expression across two groups (tumor and normal) we used the R package *DESeq2* on raw count data. The p-values were adjusted for multiple testing based on the false discovery rate (FDR) according to the Benjamini-Hochberg approach. For comparison of two patient groups two-sided Student’s t-test was used where stated, otherwise the non-parametric two-sided Wilcoxon-rank sum test was used. For comparisons among multiple patient groups one-way analysis of variance (ANOVA) and Tukey’s HSD post-hoc tests were used where stated, otherwise the none parametric Kruskal-Wallis test followed by two-sided Dunn’s pairwise post hoc tests on rank sums with Benjamini-Hochberg adjustment of p-values using the R package *PMCMR* were used. Normality of the distributions was tested with Shapiro-Wilk test and for normal distributed data the variance within each group of data was estimated and tested for equality between groups by a two-sided F-test. Distributions of data are shown either as individual data points, as box-and-whisker plots, or as violin plots. Association of CGAs with CD8^+^ or CD4^+^ T-cells was done using Spearman rank correlation and p-values were adjusted according to Benjamini-Hochberg method.

Overall survival analyses were performed using the R package *survival* and the patients were dichotomized based on median expression (NES) or divided in two or more groups by specified parameters. Kaplan Meier estimator of survival was used to construct the survival curves. Logrank tests (corresponding to a two-sided z-test) were used to compare overall survival between patients in different groups and hazard ratio (HR) (95% confidence interval) was provided for comparison of two groups. P-values were adjusted for multiple testing based on the false discovery rate (FDR) according to the Benjamini-Hochberg method. Patients for each cancer were divided in two groups based on median immunophenoscore and univariate cox regression analysis were performed and illustrated as forest plot showing log_2_(HR) and 95% confidence interval. Proportional hazard assumptions were tested. Analysis and visualization of Gene Ontology terms associated to differentially expressed genes was performed with ClueGO (Bindea et al., 2009).

### TCIA database

The web application TCIA is based on the MEAN Stack, which refers to MongoDB, ExpressJS, AngularJS and NodeJS and is completely written in JavaScript. As scaffolding tool we used the AngularJS Full-Stack Generator. AngularJS in combination with Bootstrap as front-end framework uses on the server side asynchronous data access through a RESTful Node API, built with ExpressJS. The data is stored in MongoDB, which is an open-source NoSQL database that provides a dynamic schema design. The Highcharts library is used for charting and visualization of the data. TCIA is supported by JavaScript capable browsers i.e. Google Chrome, Mozilla Firefox, Safari, Microsoft Internet Explorer, Microsoft Edge.

## SUPPLEMENTAL INFORMATION

Supplemental information includes eleven figures, six tables, and one file, and can be found online at https://icbi.i-med.ac.at/TCIA/SupplementalMaterial.

## AUTHOR CONTRIBUTIONS

Z.T conceived the project. P.C. developed the GSEA method and analyzed the data. F.F. and M.A. analyzed the neoantigens and CGAs. M.A. and M.E. estimated tumor heterogeneity and clonality of mutations. D.R. organized and managed the data transfer and storage. C.M. and D.R. developed the database. P.C. and H.H. developed the random forest approach and the immunophenogram, and analyzed the data. H.H. and Z.T interpreted the results. Z.T. wrote the manuscript.

## ACKNOWLEDGMENTS

The results shown here are in part based upon data generated by the TCGA Research Network: http://cancergenome.nih.gov. This work was supported by the Austrian Science Fund (DK Molecular Cell Biology and Oncology), the Tiroler Standortagentur (Bioinformatics Tyrol), the European Commission (Horizon2020 project APERIM: Advanced bioinformatics tools for personalised cancer immunotherapy), and the Austrian National Bank (Jubiläumsfondsprojekt Nr. 16534).

